# Interneuronal mechanisms underlying a learning-induced switch in a sensory response that anticipates changes in behavioural outcomes

**DOI:** 10.1101/2020.02.12.944553

**Authors:** Zsolt Pirger, Zita László, Souvik Naskar, Michael O’Shea, Paul R. Benjamin, György Kemenes, Ildikó Kemenes

## Abstract

How an animal responds to a particular sensory stimulus will to a great extent depend on prior experience associated with that stimulus. For instance, aversive associative learning may lead to a change in the predicted outcomes, which suppresses the behavioural response to an otherwise rewarding stimulus. However, the neuronal mechanisms of how aversive learning can result in the suppression of even a vitally important innate behaviour is not well understood. Here we used the model system of *Lymnaea stagnalis* to address the question of how an anticipated aversive outcome can alter the behavioural response to a previously effective feeding stimulus. We found that aversive classical conditioning with sucrose as the CS (conditioned stimulus) and strong touch as the aversive US (unconditioned stimulus) reverses the decision so that the same salient feeding stimulus inhibits feeding, rather than activating it. Key to the understanding of the neural mechanism underlying this switch in the behavioural response is the PlB (pleural buccal) extrinsic interneuron of the feeding network whose modulatory effects on the feeding circuit inhibit feeding. After associative aversive training, PlB is excited by sucrose to reverse its effects on the feeding response. Aversive associative learning induces a persistent change in the electrical properties of PlB that is both sufficient and necessary for the switch in the behavioural output. In addition, the strong touch used as the US during the associative training protocol can also serve as a sensitizing stimulus to lead to an enhanced defensive withdrawal response to a mild touch stimulus. This non-associative effect of the strong touch is probably based on the facilitated excitatory output of a key identified interneuron of the defensive withdrawal network, PeD12.

## INTRODUCTION

The decision to respond to a sensory stimulus in a particular way depends on variables such as prior experience. Positive experiences lead to decisions to repeat the behaviour to obtain more reward such as consumption of a palatable food item, but in a challenging environment, the value of rewards can be altered. For instance, perishing of the food item devalues the reward and can even turn it aversive. To learn to alter behavioural responses to a previously positive stimulus is essential for survival and relies on mechanisms in the nervous system that govern these highly adaptive decision making processes. Dysfunctions of these mechanisms contribute to psychiatric disorders such as addiction and depression. For example, a robust reversal of palatability (from rewarding to aversive) is observed when rats learned that a sweet tastant predicts drug induced sickness but it does not affect their self-induced cocaine administration [1]; in fact it leads to an aversive “cocaine-need state” that promotes cocaine-seeking behaviour. This and several other examples in mammals and human subjects demonstrate that learning to alter behavioural response according to a change in the value of anticipated reward is crucial for survival [2, 3]. However, the neuronal mechanisms of how this type of learning can alter the decision to perform a certain behaviour are not well understood. Activity in populations of Nucleus Accumbens (NAc) neurons in the rat is known to be related to responses to palatable (e.g. sucrose) and unpalatable (e.g. quinine) taste stimuli resulting in transiently reduced or increased firing rates respectively. Following aversive learning the reduction in response to sucrose was related to increased excitation in the NAc similarly to aversive stimulation [1]. However, the heterogeneity of neurons even within the same defined brain area, such as the NAc, makes it problematic to identify changes at the level of individual neurons. This is possible in organisms with simpler nervous systems such as *C. elegans* where a single olfactory neuron can switch between attractive and avoidance behaviors. Starving animals for as little as ten minutes in the presence of NaCl, which is normally an attractive taste, leads to salt avoidance, a reversal of normal behavior [4]. Recent findigs suggest that two components of the behavioral switch are axonal cGMP signaling and a DAG/PKC signaling pathway that affects synaptic transmission [5]. These examples indicate that learning induced alterations in behavioural response operate at multiple levels from very simple organisms such as the *C. elegans* to humans providing evidence for the existence of conserved cellular and circuit level mechanisms.

Here we used the model learning system of *Lymnaea stagnalis* [6, 7] to get further insight into the neural mechanisms of how anticipated aversive outcomes can alter the behavioural response to a previously salient feeding stimulus at the level of key identified neurons activating two opposing circuits, feeding and withdrawal. We found that applying an aversive conditioning paradigm reverses the decision so that the same stimulus inhibits feeding, rather than activating it. By pairing a food stimulus (sucrose), the CS (conditioned stimulus), with touch, an aversive US (unconditioned stimulus), the feeding stimulatory response to sucrose was reversed to an inhibition of feeding behaviour. Previous work [8] showed that aversive touch stimulus activates the defensive whole-body withdrawal circuit and simultaneously suppresses on-going feeding ingestion behaviour, even in the presence of food. We discovered that the two autonomous circuits underlying the execution of these two opposing behaviours are connected via a single type of interneuron, the PlB (pleuro-buccal) cell. PlB is an extrinsic interneuron of the feeding network that is capable of completely inhibiting feeding. It is excited by touch via a monosynaptic input from a key interneuron of the withdrawal circuit. Taking advantage of our extensive knowledge of both networks [9, 10], we now show that aversive associative learning induces a persistent change in the electrical properties of PlB that is both sufficient and necessary for the switch in the behavioural output.

## RESULTS

### Suppression of the behavioural feeding response to sucrose after aversive conditioning

Previously, we showed in *Lymnaea* that touch-induced whole-body withdrawal behaviour takes precedence over feeding and activates the neuronal mechanisms that mediate the interactions between these two incompatible hierarchically-organized behaviours [8]. Here, we tested the hypothesis that not only can aversive tactile stimulation interrupt on-going unconditioned feeding but when paired with a feeding stimulus it also leads to the conditioned suppression of the feeding response by re-programming the underlying defined circuit.

To test this hypothesis, we developed a novel type of paradigm, in which sucrose, a highly salient feeding stimulus, was used as the conditioned stimulus (CS), and aversive tactile stimulation was used as the unconditioned stimulus (US). Three groups of animals were used in this experiment (Fig. 1a). The first group (n = 12) received 5 pairings of the sucrose CS and the aversive tactile US, separated by 1 h inter-trial intervals (CS+US paired group). The second group (n = 12) also received 5 conditioning and 5 unconditioned stimuli but in an explicitly unpaired manner (10 min inter-stimulus interval), again separated by 1 h inter-trial intervals (CS+US unpaired group). Animals in both of these groups were tested for their feeding response to the CS 1 h before and again at 24 h after the first paired or unpaired trial. In the 24-h test, a third, naïve, group of animals (n = 24) was also used for comparisons with the feeding response rates in the two other groups.

**Figure 1.**
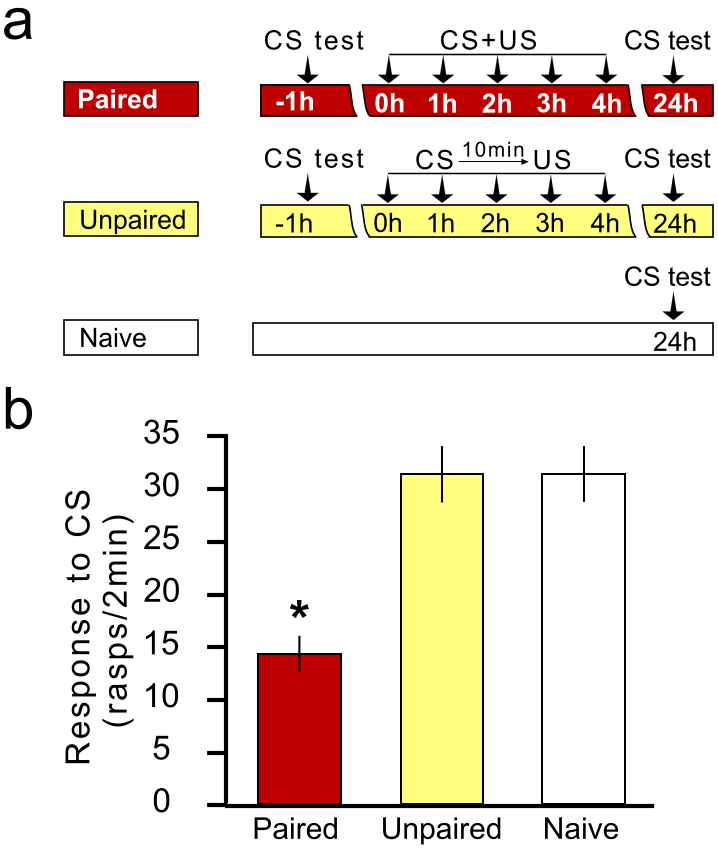
Aversive classical conditioning reduces the behavioural response to a food stimulus. a. The experimental groups and protocols used in the study. The CS (conditioned stimulus) was sucrose, the US (unconditioned stimulus) was a strong tactile stimulus to the head of the animals that evoked whole-body withdrawal. b. After aversive classical conditioning, the CS + US paired group shows a significantly inhibited feeding response to sucrose, a highly salient food stimulus in control animals. Graphs show means ± standard error of means (SEM). Asterisk indicates significance at p < 0.05.

When tested with the sucrose CS at 24 h post-training, the CS+US paired group of animals showed a significantly reduced feeding response compared to both the unpaired group and the naïve group (Fig. 1b, ANOVA p < 0.0001, Tukey’s tests: CS+US paired group versus CS+US unpaired and Naïve group, p < 0.05 both; CS+US unpaired group versus Naïve, p > 0.05), and also when compared to the mean pre-training response levels to the CS in the same group (35.3 ± 3.9 rasps/2 min, paired t-test: p < 0.0001). This reduction in the feeding response was specific to the sucrose stimulus; when another salient feeding stimulus, fresh cucumber juice was applied 24 h after aversive training with sucrose as the CS, the animals (n = 12) showed the same high levels of feeding response as to sucrose before training (Supplemental Fig. 1, paired t-test: p = 0.4) and as the unpaired and naïve control animals after training (ANOVA: p = 0.98). These behavioural experiments therefore established that when sucrose is paired with aversive tactile stimulation, an association is formed between the two stimuli, which results in a selective impairment of the feeding response to sucrose without affecting the functioning of the feeding network in general.

While observing the feeding response to the sucrose CS 24 hours after paired or unpaired training and also in naïve animals, we did not see any withdrawal reactions in response to the CS in any of the groups. This indicated that unlike the pairing of a non-food chemosensory conditioned stimulus with an aversive unconditioned stimulus in *Aplysia* [11], the pairing of a salient food chemosensory conditioned stimulus with an aversive unconditioned stimulus in *Lymnaea* does not lead to conditioned fear.

### Interneuronal mechanism of feeding inhibition after aversive conditioning

We reasoned that a long-term reversal of the inhibitory response of the PlB interneurons to sucrose-activated sensory inputs underlies the observed long-term behavioural changes induced by aversive conditioning.

To test this hypothesis, we compared the response of PlB neurons to sucrose in semi-intact lip-CNS preparations made from snails that had been subjected either to the paired or unpaired protocol or were from naïve animals (n = 10 in each group). In some of these preparations (paired, n = 3; unpaired, n = 2; naïve, n = 3) we also left the buccal mass, the main effector organ for feeding [12], attached to the CNS, which allowed the monitoring of spontaneous and sucrose evoked feeding behaviour by measuring its rhythmic contraction activity. In both naïve and unpaired animals, sucrose application to the lips hyperpolarized PlB and reduced its tonic firing and this was accompanied by an increase in the frequency of muscular contractions of the buccal mass (Fig. 2a, b), indicating that sucrose is stimulating ingestive feeding behaviour. By contrast, in preparations from aversively trained animals the PlB cell showed both a marked depolarization and a consequent increase in its tonic firing rate in response to the application of sucrose to the lips (Fig. 2c). These responses were accompanied by a cessation of rhythmic buccal mass contractions underlying spontaneous feeding (Fig. 2c). Spontaneous buccal mass contractions were observed in all three lip-CNS-buccal mass preparations from aversively trained animals (example in Fig. 2c) whereas the same type of preparations made from snails that had been subjected either to the unpaired protocol or were from naïve animals showed no or only occasional contractions of the buccal mass (examples in Figs. 2a and b).

**Figure 2.**
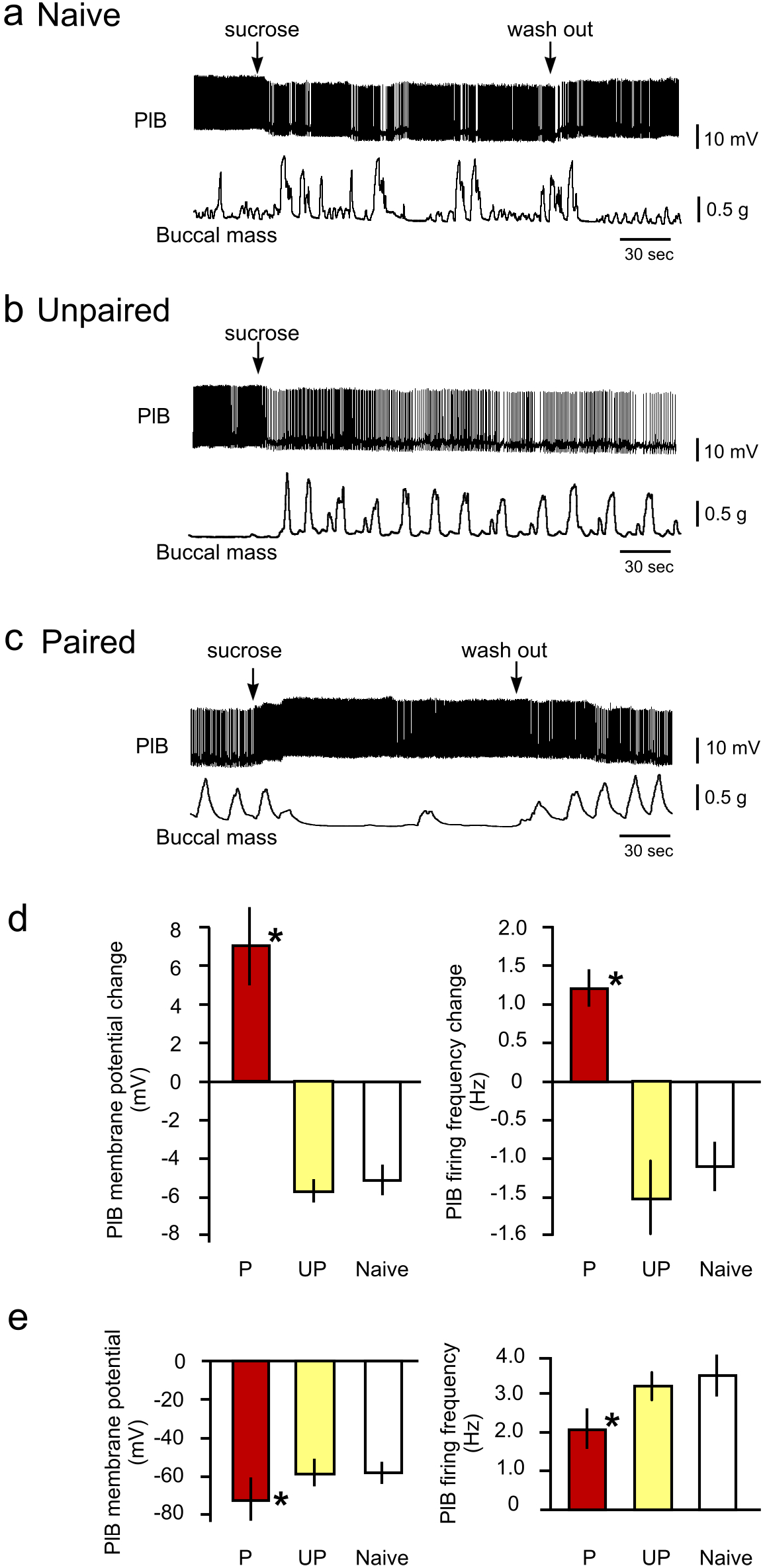
Aversive classical conditioning reverses the electrophysiological response of an identified modulatory interneuron to a food stimulus. a, b. In typical semi-intact preparations from naïve and unpaired control animals, the application of sucrose to the lips results in the hyperpolarization and resulting reduction of the firing rate of the modulatory interneuron PlB and the triggering of rhythmic contractions of the buccal mass. The Unpaired trace in b illustrates that if the sucrose stimulus is not removed from the sensory areas, PlB remains hyperpolarized with a correspondingly lower spike frequency, and rhythmic feeding activity continues. c. Representative example of a semi-intact preparation from a CS + US paired animal where the application of sucrose to the lips results in the depolarization and resulting increase of the firing rate of PlB and the cessation of the spontaneous rhythmic contractions of the buccal mass. d. Statistical comparisons of the sucrose-evoked electrophysiological responses of PlBs in preparations from CS + US paired (P) and control (unpaired (UP) and naïve) animals. e. Statistical comparisons of intrinsic electrical properties of PlBs in preparations from CS + US paired (P) and control (unpaired (UP) and naïve) animals. Graphs in d and e show means ± standard error of means (SEM). Asterisks indicate statistical significance of at least p < 0.05.

When compared among the naïve (n = 10), unpaired (n = 10) and paired groups (n = 10), both the changes in PlB membrane potential and firing rate in response to sucrose were consistently in the opposite direction and highly significantly different in the paired versus the other two groups (Fig. 2d, ANOVA: p < 0.0003, Tukey’s: p < 0.001 for both parameters), while not different in the naïve and unpaired groups (Tukey’s: p > 0.05). This result shows that by increasing PlB’s firing rate sucrose is now strongly activating the inhibitory PlB to feeding circuit pathway known to be involved in the suppression rather than the activation of feeding [13].

An analysis of the same experiments also revealed a significant difference between both the baseline membrane potential and spike frequency levels of PlBs in preparations from paired versus both naïve and unpaired control animals, when measured 24 h after training (Fig. 2e). Specifically, PlBs from CS+US trained animals had a significantly more hyperpolarized membrane potential (−68.5 ± 4.8 mV) and lower baseline spike frequency (2.0 ± 0.3 Hz) compared to PlBs from naïve animals (−57.4 ±1.5 mV and 3.3 ± 0.4 Hz, respectively) and unpaired animals (−58.1 ± 2.0 and 3.1 ± 0.2 Hz, respectively) (Dunnett’s tests, p < 0.05). These experiments therefore showed that aversive conditioning changed key intrinsic cellular properties of the PlBs.

### Photo-inactivation of a single PlB reverses the effects of aversive conditioning on sensory inhibition

To test if PlB was necessary for the suppression of sucrose-driven feeding response after aversive training, we examined the effects of PlB photoinactivation on the feeding motoneuronal responses to sucrose. Intracellular recording from the PlB neuron shows that the resting membrane potential changes from −53 ± 8 mV to 0 mV within 7 minutes from the beginning of photoinactivation (Fig. 3a, b) leading to the cessation of spike activity. Changes in spontaneous rhythmic motoneuronal (B3) feeding activity following application of sucrose were compared before and after the photoinactivation of single PlB neurons (Fig. 3c, d). Prior to PlB photoinactivation, in preparations from animals subjected to paired training sucrose caused the cessation of spontaneous fictive feeding activity and failed to trigger rhythmic food-induced activity, whereas after photo-inactivation, the same preparations regained their ability to respond to the sucrose stimulus with increased fictive feeding (Fig. 3c). A comparison of the fictive feeding cycles recorded in B3 after the application of sucrose showed a significant difference between the feeding rates before and after PlB had been photo-inactivated (n = 8 preparations, paired t-test: p < 0.0003). These experiments demonstrated that inactivation of a single PlB removes the effects of aversive conditioning providing strong evidence that the PlB interneurons are the main locus for the learning-induced changes in the sucrose response. The photoinactivation of only one of the paired PlB cells is sufficient to remove the effects of aversive conditioning.

**Figure 3.**
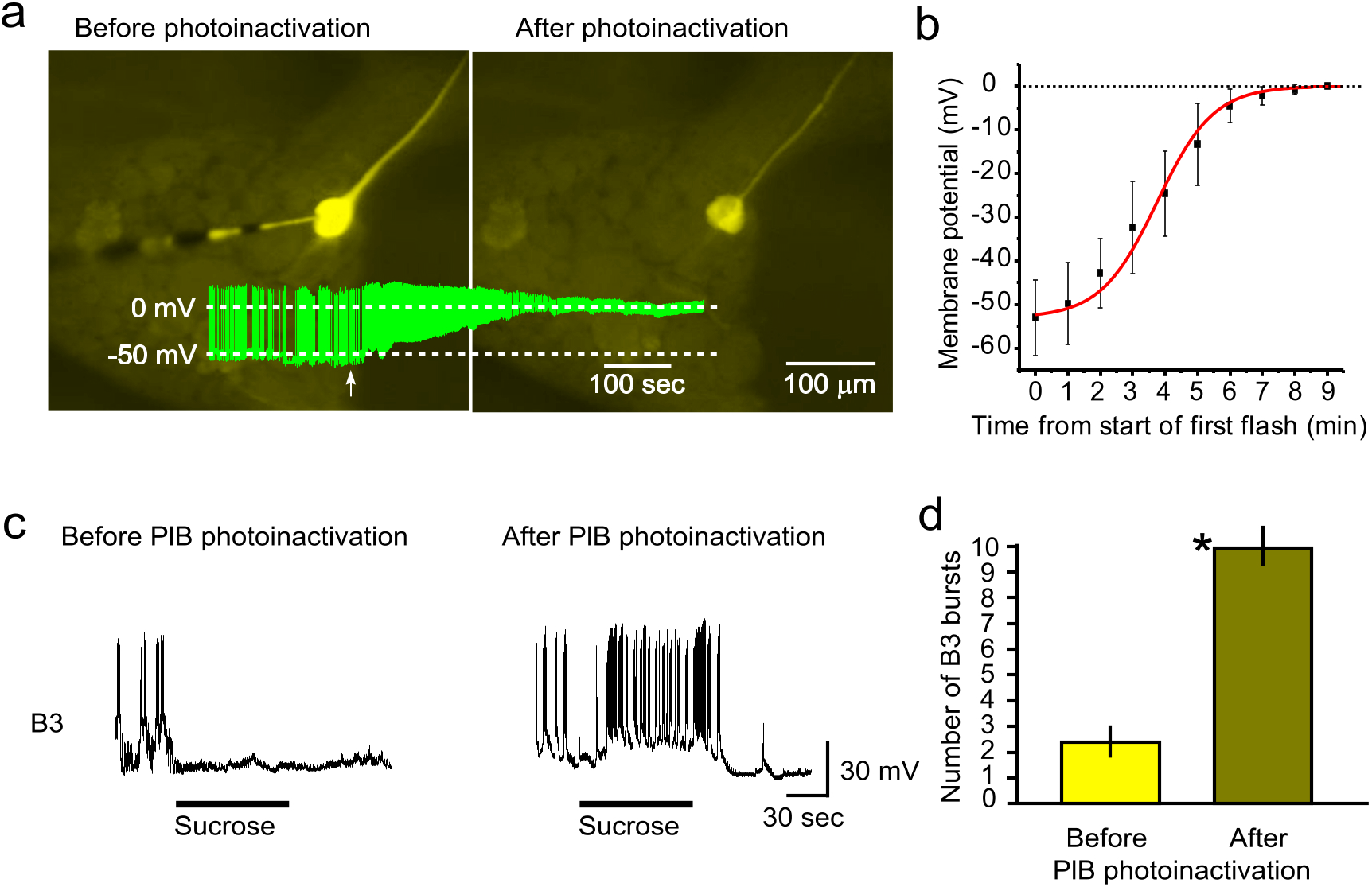
Photoinactivation of PlB converts the paired-preparation phenotype to a naïve one. a, b. The temporal progression of the loss of the electrical properties (firing and membrane potential) of a carboxyfluorescein-filled PlB neuron by photoinactivation. c. Left, cessation of spontaneous and absence of sucrose-induced fictive feeding cycles recorded on a B3 feeding motoneuron in a semi-intact preparation from a CS + US paired animal before PlB photoinactivation. Right, After PlB photoinactivation in the same preparation, the feeding motoneuron shows fictive feeding cycles in response to sucrose applied to the lips. d. Statistical comparison of the number of fictive feeding cycles recorded in B3 during the application of sucrose. Graphs show means ± standard error of means (SEM). Asterisk indicates significance at p < 0.05.

### The withdrawal response interneuron PeD12 is not involved in inhibitory sucrose responses following aversive conditioning

Our previous study [8] showed that the withdrawal response interneuron, PeD12, is the key element in touch-induced inhibition of feeding. Touch excites PeD12 and via a monosynaptic excitatory pathway increases the tonic firing of the PlB leading to the suppression of feeding. Could the conditioned aversive effect of sucrose also be mediated by this interneuronal pathway? To test this hypothesis, we set up semi-intact lip-CNS preparations from CS+US trained, unpaired and naïve control animals and co-recorded PeD12 with a feeding motoneuron, B4, to establish if PeD12 responds to the sucrose CS after paired training. We found that although the fictive feeding response to sucrose was suppressed, PeD12 did not respond to the sucrose CS after aversive learning (Fig. 4) and so did not to play a role in the blocking of feeding. This finding provides evidence that the post-training aversive effect of sucrose is mediated by a different pathway than the one that is activated by a strong aversive touch.

**Figure 4.**
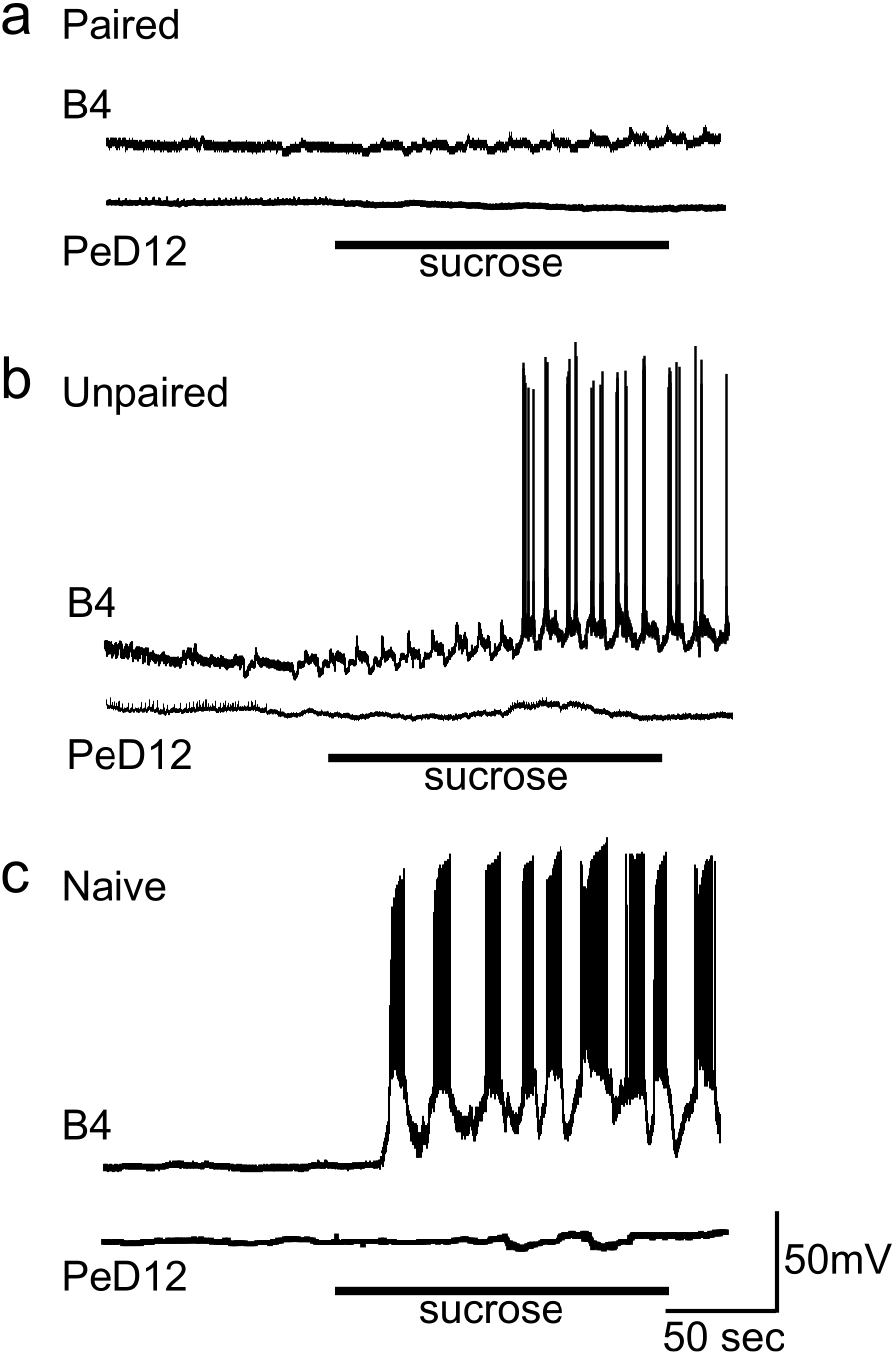
The conditioned aversive effect of sucrose applied to the lips in semi-intact preparations is not mediated by the withdrawal interneuron PeD12. a. Simultaneous recordings of the feeding motoneuron B4 and the withdrawal interneuron PeD12 in a preparation made from a CS + US paired animal showing the lack of a fictive feeding response in B4, in the absence of sucrose-triggered activation of PeD12. b, c. Sucrose activates fictive feeding in preparations from an unpaired and a naïve animal, respectively. As expected, the withdrawal interneuron, PeD12, shows no activation from the application of a rewarding food stimulus, sucrose.

### Aversive training facilitates the PeD12 to PlB synaptic connection and results in sensitized avoidance response to weak touch

Even though PeD12 did not respond to the aversively conditioned sucrose stimulus, we found that 24 h after training an artificially triggered burst of PeD12 action potentials resulted in a significant increase in the postsynaptic depolarization and spike frequency recorded in the PlB in preparations from CS+US paired (n = 17) as well as unpaired animals (n =13) compared to naïve controls (n = 18) (Kruskall-Wallis tests: p < 0.003 (EPSP size), p < 0.0001 (spike frequency); Dunn’s multiple comparisons tests: p < 0.05 for both Naïve versus Paired and Naïve versus Unpaired for both EPSP size and spike frequency, Fig. 5). This observation indicated that the excitatory synaptic output of PeD12 becomes facilitated when a strong tactile stimulus is applied to intact *Lymnaea*. This led us to setting up the general hypothesis that the aversively conditioned animals as well as undergoing associative behavioural plasticity might become sensitized by the strong tactile stimuli, which are known to trigger bursts of action potentials in PeD12 leading to whole-body withdrawal [8].

**Figure 5.**
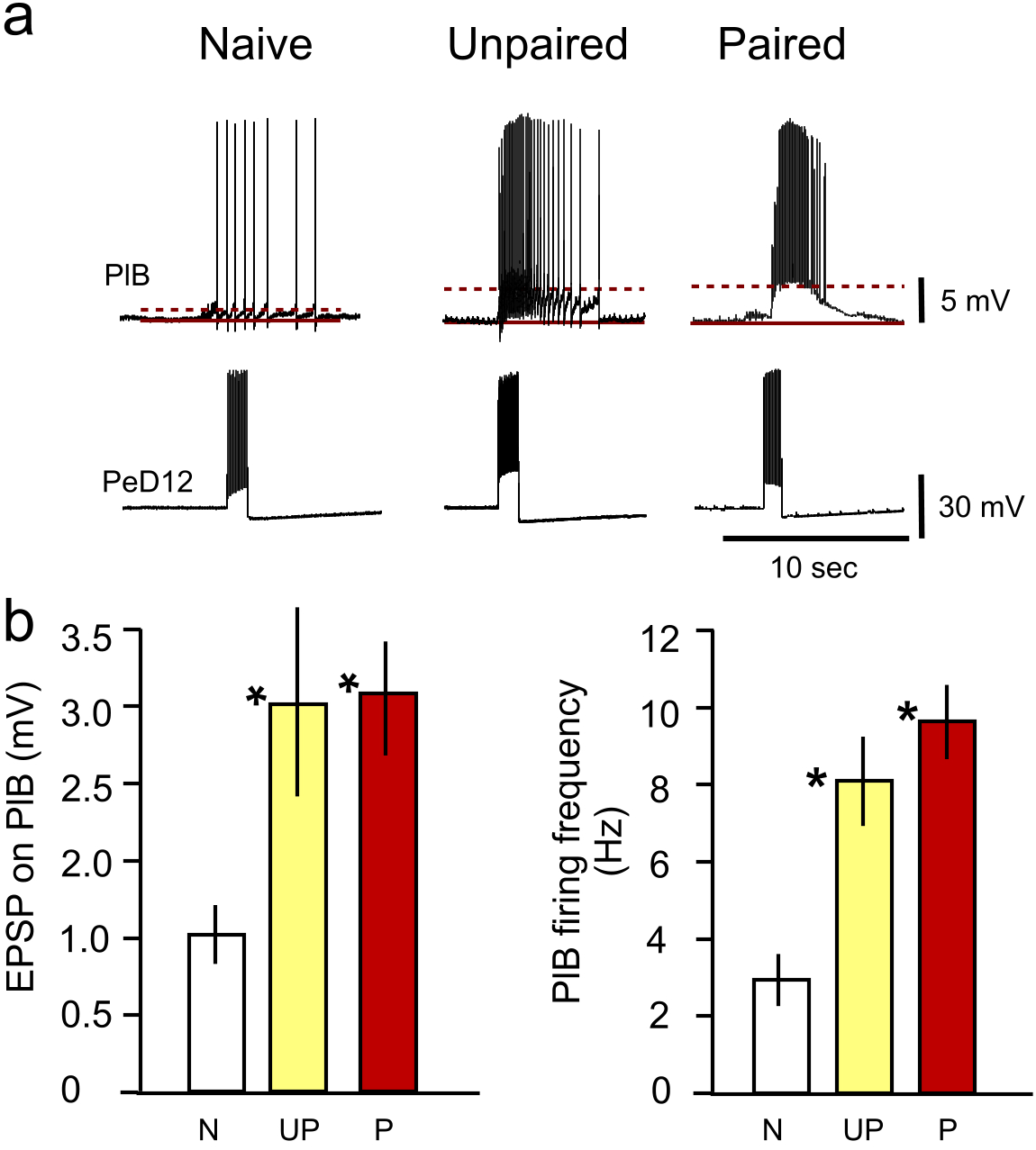
Long-term facilitation of the PeD12 to PlB synapse after paired and unpaired training using the same strong aversive tactile stimulus. a. Square-pulse triggered bursts of action potentials in PeD12 evoke facilitated excitatory postsynaptic responses in preparations from both a CS + US paired and an unpaired animal compared to a preparation from a naïve animal. b. Statistical comparisons of the PeD12-triggered PlB EPSP peak amplitudes in preparations from naïve (N) versus unpaired (UP) and CS + US paired (P) animals. Graphs show means ± standard error of means (SEM). Asterisks indicate significance of at least p < 0.05 (compared against the naïve data).

The specific hypothesis to be tested was that a weak tactile stimulus that normally does not evoke a strong aversive response [14] would do so if a strong tactile stimulus was applied in the same temporal pattern as the aversive conditioning but without the sucrose CS. Aversive responses to the week tactile stimulus were measured 24 hours later and compared with pre-training behavioural responses to the same weak tactile stimulus using visual analysis of video clips that, unknown to the observer, were from either pre or post training animals. One observer achieved a perfect match (25/25, 100%) to the real identity of the video clips (pre- or post-training test) while the other three had a 24/25 (96%) success rate each, all significantly above chance (50%) (two-tailed one-sample z-test, p<0.05), indicating that the snails become sensitized to the weak touch stimulus.

## DISCUSSION

Sensory cues in the natural environment predict reward or punishment and rapid learning of the meaning of such indicators is vital for the survival of the organism. For example, the ability to detect attractive tastes or odours such as palatable food, or potential mates is essential for foraging and social behaviour while recognition of harmful or inedible substrates and predators are crucial for avoidance of malaise and escape from harm. While some of these reactions are innate like the stereotyped avoidance behaviour to volatile odours (alcohols, ketones) and carbon dioxide in *C. elegans* [15, 16], others are learned, for example the avoidance of some pathogen bacteria through association with illness [3]. The main goal of our study was to elucidate how associative learning can modify a strong innate behaviour, the response to food, taking advantage of an invertebrate model organism, *Lymnaea*, which has well-defined neuronal circuits underlying both feeding and withdrawal escape behaviours [10, 17, 18]. Sucrose is a salient appetitive stimulus that evokes vigorous feeding response in many animal species, including *Lymnaea* [19, 20]. Touch to the head of the animal initiates a withdrawal reflex that disrupts ongoing feeding responses to sucrose. Escape is higher in the behavioural hierarchy than feeding [8] and therefore overrides the innate urge to feed. Random application of head touch does not lead to long-lasting alteration of the innate feeding response. However, we show here that aversive classical conditioning using touch [12] leads to a subsequent impairment of the feeding, reducing the sensory response to sucrose, the innate highly salient food stimulus. Previously, we have shown that in isolated brain preparations from animals aversively conditioned with L-serine, an appetitive chemical CS and quinine, an aversive chemical US, an *in vitro* analogue of the conditioned stimulus (electrical stimulation of the median lip nerve) caused a significant increase in PlB firing rate compared with naïve controls as well as a lower expression of fictive feeding cycles (an *in vitro* correlate of the conditioned response) [21]. Although this earlier work identified PlB as a neuronal substrate for aversive learning in general, it did not link it directly to its already known role in triggering the inhibition of feeding in response to strong tactile stimulation [8]. In the present study the use of an aversive paradigm where the US was a strong touch has now allowed us to make this link by demonstrating that in semi-intact preparations from aversively conditioned animals the application of sucrose (the CS) to the lips increases PlB firing frequency and inhibits feeding, whereas in control animals that received unpaired training or no training the opposite happened: PlB firing frequency decreased and feeding was activated in response to the sucrose stimulus.

Studies in *C. elegans* showed that there are two types of sensory neurons (AWA and AWC) responsible for odor attraction responses while a single neuron type (AWB) directs odour avoidance [15] [22]. Interestingly, genetic manipulation of these neurons reveals that the modification of the presynaptic pathway in AWC axons can switch the behavioural response indicating that the intrinsic property, such as its connectivity or the synaptic release properties of the neuron guiding the odour response, is independent of the responding receptor [5].

Our findings indicate that similar to *C. elegans*, *Lymnaea* also has a single cell type that is crucial for the switching between avoidance (whole body withdrawal) and attraction (feeding). However, in the case of *Lymnaea*, the decision is made by an interneuron (PlB) rather than at the level of sensory neurons and importantly, it can be altered by learning. Simultaneous recordings of intracellular activity from identified neurons of both the feeding and the withdrawal circuits revealed that by increasing the firing frequency of the PlB neuron by positive current injection the feeding network can be supressed whilst reducing firing by hyperpolarization leads to activation of feeding [23].

Importantly, by using semi-intact lip-brain preparations we also found that after aversive learning, the firing frequency of the PlB increases rather than decreases in response to application of sucrose to the lips. This shows that after conditioning, this interneuron is capable of suppressing a strong innate behavioural response. Similar changes were observed in vertebrates where populations of inhibitory neurons in the NAc increased their firing rate after aversive conditioning causing the suppression of the feeding response [1].

Extensive work on insects concentrated on identifying the molecular and cellular mechanisms of odour aversion and attraction in pursuit of finding successful insect repellents and traps in order to combat disease and agricultural pests. These studies also highlighted that neurons with related sensory receptors and similar projections to the olfactory bulb can generate opposing behaviours. Although it is difficult to identify individual neurons in the insect brain, using ablations in *Drosophila* it was found that there are two higher order olfactory nuclei, the lateral horn and the mushroom body, which are thought to mediate innate and learnt odour responses, respectively [24–26].

In *Lymnaea*, a single interneuron (PlB) mediates the learnt response that alters innate feeding behaviour. Although this neuron is extrinsic to both the feeding and the withdrawal circuits, it can supress the feeding response when the withdrawal network is activated. As we have shown previously (8), this is achieved via an excitatory monosynaptic connection from one of the neurons (PeD12) of the withdrawal circuit. However, our finding that PeD12 is not activated by sucrose in preparations from aversively trained animals shows that the post-training aversive effect of sucrose is mediated by a pathway that is different from the pathway activated by a strong aversive touch. However, both these pathways share PlB as their final target. This interneuron provides the inhibitory input to the feeding network necessary for the cessation of feeding both in response to an unconditioned mechanosensory stimulus [8] and a conditioned sucrose stimulus.

Although our results showed that PlB is necessary for the generation of the conditioned behavioural response, we still lacked an explanation for the mechanism underlying the long-term change in this interneuron’s firing properties. Intracellular recording of electrical parameters from PlBs in trained and control preparations provided the answer to this question. Notably, the resting membrane potential in trained PlBs was significantly more hyperpolarized than in PlBs from both unpaired and naïve control animals. This was accompanied by a significantly lower rate of spontaneous tonic spike activity in PlBs recorded in preparations from paired versus both naïve and unpaired animals, which may explain the apparently higher spontaneous feeding rate see in the lip-CNS-buccal mass preparations from animals that had been subjected to aversive conditioning. These important observations, together with the finding that after aversive training, PeD12 is not activated by the CS, lend strong support to the conclusion that aversive training causes persistent changes in key intrinsic properties of PlB, which underpin its fundamentally altered response to food.

Our previous work has shown that after appetitive conditioning using either a tactile or a chemical CS and a sucrose US, extrinsic modulatory interneurons of the feeding system undergo a persistent depolarization that in turn allows the gating in of the circuit response to the CS in trained animals [27, 28]. Based on these previous findings, we would have expected PlB to become more rather than less depolarized after aversive learning. However, this was not the case: PlB became persistently hyperpolarized in aversively conditioned animals. The baseline electrical properties of the PlB in naïve animals may provide an explanation for why the learning-induced intrinsic changes in PlB are different from those in the other two previously studied extrinsic interneurons of the feeding system, the command-like Cerebral-Ventral A (CV1a) cell and the gating/modulatory interneuron Cerebral Giant Cell (CGC) [9]. Following appetitive conditioning, a persistent non-synaptic change (long-term depolarization) occurs in one or other of these neurons depending on the type of conditioning [28, 29]. Studies using conditioned taste aversion (CTA) rather than tactile-based aversive training shows that ablation of the CGC cell bodies before, but not after training, disrupted memory formation [30]. Furthermore, unlike the appetitive chemical training, CTA does not result in changes of electrical properties of these important interneurons of the feeding system [31]. Employing a tactile (touch to the head) stimulus as an aversive US, just like in the present study, results in no significant change in the activity of the CGCs [32].

Unlike either CV1a or CGC, the PlB fires tonically at a very high frequency at its baseline activity level in preparations from both naïve and unpaired control animals. Food-triggered sensory input hyperpolarizes PlB and reduces its firing frequency, which reduces its overall inhibition of the feeding network. The likely reason for the observed training-induced persistent baseline hyperpolarization and reduction in spike frequency is that these allow the de-inactivation of ion channels that are involved in driving the strong depolarization and resulting significant increase in spike frequency necessary for PlB to increase its firing in response to the CS that underlies the suppression of the feeding response.

The same strong tactile stimulation that served as the US in the aversive conditioning experiments may also serve as a sensitizing stimulus leading to a persistently sensitized withdrawal response to mild tactile stimulation. The monosynaptic excitatory input to PlB from PeD12, a key interneuron of the withdrawal network [8], was facilitated in preparations from both the paired and unpaired groups of animals, which suggested that it was due to a non-associative effect of the training paradigms used in our experiment.

Although PlB is not a member of the withdrawal network, our previous work demonstrated that this neuron is activated by PeD12 when a strong tactile stimulus is applied to the lips of a semi-intact preparation, and this leads to the termination of on-going feeding while the PeD12-driven activation of the withdrawal circuit drives the whole-body withdrawal response [8]. It was therefore reasonable to assume that if the PeD12 to PlB connection becomes facilitated by the strong tactile stimulus, so do the previously described [8] PeD12 to withdrawal circuit neuron connections, resulting in a sensitized withdrawal response to mild touch. So next, we tested the hypothesis that in intact animals the application of a strong tactile stimulus alone can serve as sensitization training. This hypothesis was validated by the finding that animals exposed to sensitization training showed an enhanced withdrawal response to a mild touch 24 hours later. This suggests that in our aversive conditioning experiments, where animals had been given strong tactile stimuli during training, the strong conditioned suppression of feeding behaviour emerged in parallel with a sensitized defensive response, underpinned by facilitated PeD12 presynaptic activity.

Interestingly, when sensitization training with strong electric shocks to the body wall was used in *Aplysia*, it resulted in a persistent suppression of feeding behaviour as well as a sensitized defensive response to a brief, mild current pulse to the tail [33]. Based on these previous findings we would have expected to see a suppressed feeding response in both the Paired and Unpaired groups of animals in our experiments. However, we only found a reduced feeding response after paired training indicating that aversive associative and non-associative training may have different effects on the neural circuits controlling the withdrawal and feeding network.

Our findings also have some interesting similarities to early seminal findings in *Aplysia* that provided the first evidence for conditioned fear in an invertebrate [11]. In that study, an initially neutral chemosensory CS was paired with electric shock to the head as the US. This aversive classical conditioning resulted both in the facilitation of a number of defensive responses to the CS and a concomitant depression of the feeding response to seaweed, a natural salient food stimulus for *Aplysia* [11]. However, our aversive classical conditioning paradigm was based on the use of a salient food chemosensory CS, and its primary effect was a pairing-specific depression of the feeding response to this CS, in the absence of a defensive withdrawal response to it. At the same time, the strong tactile US used in our experiments caused a sensitized response to a mild tactile stimulus at test. Therefore, unlike the paradigm used in the *Aplysia* study, which resulted in a variety of different conditioned fear responses to the CS, including feeding depression [11], our paradigm seems to have resulted in a conditioned suppression of feeding and a non-associative enhancement of the withdrawal response without the emergence of conditioned fear. From this we conclude that unlike the noxious electrical stimulation used in *Aplysia* [11], the aversive tactile stimulation used in our experiments did not produce defensive arousal.

Our new results provide evidence for a neural mechanism underlying the switch from a very high to a low punishment error prediction after aversive associative learning based on the use of a very salient feeding stimulus as the CS. This results in the suppression of a strong innate behavioural response according to a learned change in the value of anticipated reward. Another major finding is that a non-associative learning mechanism, sensitization, also plays a role by changing another behavioural response, withdrawal to a mild touch, through an alternative neural pathway.

## ACKNOWLEDGEMENTS

This work was funded by the Biology and Biotechnology Research Council grants BBSRC/BB/H009906/1 (M.O.S., G.K., P.R.B.) and BBSRC/BB/P00766X/1 (I.K., G.K., P.R.B.) and the National Brain Project (Hungary) grant 2017-1.2.1-NKP-2017-00002 (Z.P.).

## AUTHOR CONTRIBUTIONS

Z.P., M.O.S., P.R.B., G.K. and I.K. conceived and designed the experiments. Z.P., Z.L. and S.N. performed the experiments. Z.P., G.K. and I.K. analyzed the data and made figures. Z.P., P.R.B., G.K. and I.K. wrote the paper. All the authors read and contributed to the submitted version of the manuscript. M.O.S., I.K., G.K. and P.R.B. acquired the funding and were responsible for resources.

## COMPETING INTERESTS

The authors declare no competing interests.

## MATERIALS & CORRESPONDENCE

All correspondence and material requests should be addressed to.

## EXPERIMENTAL PROCEDURES

### Experimental animals and behavioural procedures

#### Animals

We used *Lymnaea stagnalis* from a laboratory-bred stock of adult (5-6 months old) snails. Animals were kept in groups in large holding tanks containing copper-free water at 20-21°C on a 12:12h light-dark regime. The animals were fed lettuce three times and a vegetable based fish food (Tetra-Phyll; TETRA Werke, Melle, Germany) twice a week, except before starting an experiment, when they were food-deprived for two days.

#### Multiple-trial aversive classical conditioning

Snails were trained using a multiple-trial aversive classical conditioning protocol in which the order of the two stimuli (touch and sucrose) originally used for reward conditioning [14, 23, 34, 35] was reversed and the concentration of sucrose was reduced while the intensity and number of tactile stimuli used in each trial was increased. Thus in this paradigm, in each trial 6.7% sucrose (starting concentration in 5 ml copper-free water), the conditioning stimulus (CS), was paired with a series of aversive tactile stimuli as the unconditioned stimulus (US).

Before aversive training, the snails were placed individually into Petri dishes containing 95 ml of copper-free water for a 10 min acclimatization period, so that a low level of spontaneous rasping was reached in the novel environment [36]. During each trial of the tactile classical conditioning protocol, the snails were first presented with the CS (final concentration 0.34% in 100 ml copper-free water). After 15 sec (when all the freely moving snails had started feeding), the US was presented using a hand-held probe with a tip made of a tooth-pick. The target zone on the lip structure was the median portion adjacent to the mouthparts including the leading edge of the lips as previously described by Staras et al. [19]. During the rest of the 2 min training period, the procedure was repeated every 15 sec. The pairing of sucrose with a series of 7 tactile stimuli constituted one trial. After this, the animals were rinsed in a clean water tank to remove any residual sucrose before they were placed back into their home tank. Sixty minutes after the first trial, the animals received a second training trial followed by a third, fourth and fifth trial another 60 min later (Figure 1a). For explicitly unpaired control, snails received five presentations of the CS and US with 10 min inter-stimulus intervals, in contrast to the 15 sec used in the paired training protocol. Similar to the paired training protocol, snails were placed individually into Petri dishes for a 10 min acclimatization period. The snails were presented with the CS in the 0.34% final concentration without US. After a 2 min period, the animals were transferred to another Petri dish (100 ml copper-free water) for 10 min, after which the US was presented 7 times with 15 sec intervals for 2 min in each trial. Thereafter, the animals were put back into their home tank. Sixty minutes after the first trial, the animals received the rest of the 4 training trials, again at 1 h intervals. CS tests were performed both before and after paired and unpaired training (Figure 1a).

Twenty-four hours after the first trial, individual snails were taken from their home tanks using a blind procedure and placed in Petri dishes for testing the response to the CS. After a 10 min acclimatization period, rasps were counted for 2 min (i.e., spontaneous rasping in water). Sucrose was then applied and rasps were counted for a further 2 min (i.e., the feeding response to the sucrose CS). For testing the integrity of the feeding network of trained animals as well as the stimulus specificity of the conditioned response after aversive training with sucrose and strong touch, an additional control experiment was performed. In this experiment the number of rasps was counted after presenting filtered cucumber juice to the animals as a feeding stimulus other than sucrose.

A third, naïve control group of animals, was also used in the experiments. These animals were kept in the same conditions as the experimental ones but only tested once with the CS at the same time when the experimental groups were tested 24 h after the paired or unpaired training.

#### Sensitization training

The pre-training treatment of the snails (n = 25) with the CS in this experiment was the same as described for the aversive conditioning experiment. However, in a pre-training test, all 25 animals were presented with a weak tactile stimulus to the head using a hand-held probe with a tip made of a thin wedge of soft, flexible plastic [34], while their behaviour was recorded by the video camera of an Apple iPhone 6S+.

During the 2-min training periods, the same strong tactile stimulation to the lip region that was used as the US in the aversive training protocol, was repeated every 15 s in each trial but in the absence of the sucrose CS. After this, the animals were placed back into the home tank. Sixty minutes after the first trial, the animals received a second US only training trial followed by a third, fourth and fifth trials another 60 min later.

After 24 h, individual snails were taken for testing from their home tanks and placed in Petri dishes. After a 10-min acclimatization period, the same weak tactile stimulation that was used in the pre-tests was applied to the lip region of the animals while they were being videod. The pre- and post-training video clips were randomly labeled A or B and shown to 4 different members of the laboratory who had no knowledge of which label belonged to the pre-versus post-training video clips but were highly trained in observing aversive behavioral responses in *Lymnaea*. The testers had to establish which clip taken of the same individual animals showed a stronger aversive response to the same weak stimulus.

### Preparations and electrophysiological recordings

#### Preparations

Experiments were performed on semi-intact preparations containing the entire CNS and attached sensory regions (lips and tentacles) [18, 23, 34, 36]. A modified semi-intact preparation, containing the buccal mass, the main feeding muscle, was used to measure rhythmic contractions induced or inhibited by sucrose.

#### Transducer recording

Rhythmic buccal mass contractions in the preparations were activated or inhibited by sucrose (0.02 mM in normal saline) applied to the lips via a computer controlled gravity-fed perfusion system. At the systems level, the sucrose induced responses were observed both on the buccal mass and on the feeding motoneurons (B3 or B4) confirming that they were generated by the feeding network. The buccal mass was attached to a force transducer (WPI FORT10g, World Precision Instruments, Incorporation, Sarasota, USA). Muscle recordings were made by connecting the force transducer through a Digidata 1320A interface (Axon Instruments, Union City, CA, USA) to a PC.

#### Electrophysiology

Preparations were dissected and neurons recorded in a Sylgard-lined chamber containing normal snail saline (50 mM NaCl, 1.6 mM KCl, 3.5 mM CaCl_2_, 2.0 mM MgCl_2_, 10 mM HEPES, pH 7.9). The outer layer of the thick connective tissue sheath was removed mechanically from the dorsal surface of ganglia and the inner layers were softened by 1% protease treatment (Sigma XIV, Sigma) for 2 min.

Intracellular recordings were performed under a stereomicroscope (Leica MZ FLIII, Switzerland). AxoClamp 2B (Axon Instruments, Union City, CA, USA) and NeuroLog D.C. (Digitimer Ltd., UK) amplifiers were used to monitor the electrical activity of identified neurons. Membrane potential (MP) manipulation was carried out by current injection through the recording electrode. Microelectrodes were pulled from borosilicate glass pipettes (GC200F-15, Harvard Apparatus, UK) with Narishige (Narishige Scientific Instrument Laboratory, Japan) vertical puller to a 15–20 MΩ tip resistance when filled with 4 M potassium acetate. For data acquisition and protocols, the amplifiers were connected via a DigiData 1320A interface (Axon Instruments, Union City, CA, USA) to a PC supplied with pClamp8.2 software (Axon Instruments, Union City, CA, USA). The recorded traces were analysed by OriginLab Corporation Origin 8.5 software.

#### Identification of neurons

The left and right B3 feeding motoneurons of the buccal ganglia were mainly used to monitor the central pattern generator (CPG)-driven fictive feeding rhythm. In some experiments B4 feeding motoneurons of the buccal ganglia were recorded to confirm the occurrence of feeding cycles. Both types of neuron can be identified by size, location and characteristic fictive feeding activity [9, 12].

The PlB (Pleural-Buccal) neuron is an extrinsic modulatory interneuron that inhibits the feeding network [13]. It is a small neuron (20-30 μm cell body diameter) that lies on the medial surface of the pleural ganglion close to the pleural-pedal connective [13, 23]. It shows characteristic tonic firing activity and its identification is confirmed by recording inhibitory synaptic responses on feeding neurons.

PeD12 is a recently described interneuron (60 μm in cell body diameter) of the whole-body withdrawal network and its artificial stimulation excites the PlB monosynaptically causing inhibition of ongoing feeding [23]. It lies close to the previously described PeD11 [38] on the dorsal surface of the pedal ganglia close to the statocyst.

#### Intracellular dye-injection and photoinactivation

To photoinactivate the feeding inhibitory PlB cells, they were filled with the fluorescent dye Lucifer Yellow dilithium salt (10 mM, Sigma, UK). After identification of PlB, the dye was loaded into the cell bodies from the recording microelectrode by a 10 ms and 5-8 psi pulse of a multi-channel picospritzer (General Valve Corporation, New Jersey, USA). The loaded PlB cells were photoinactivated by continuous high-energy UV flash lamp pulses using a JML-C2 (Rapp OptoElectronic, Hamburg, Germany). During photoinactivation, 1000 μs length (C3 capacitor mode) and 180-200 J energy pulses were used for up to 8 min. The JML-C2 was triggered by an external TTL signal. The membrane potential of PlB was monitored continuously during the flash photolysis procedure by pClamp8.2 software (Axon Instruments, Union City, CA, USA).

#### Statistical methods

In both behavioural and *in vitro* experiments comparisons between more than two independent groups (e.g., naive, unpaired, trained) were carried out using ANOVA or Kruskall-Wallis tests followed by multiple post-hoc Tukey’s, Dunnet’s or Dunn’s tests. When comparing only two independent samples, t-tests were used. All statistical analyses were carried out using Prism (GraphPad) software. The differences were considered statistically significant at p < 0.05. For comparison against an expected single value, a two-tailed one-sample z-test was used.

#### Data availability statement

The data that support the findings of this study are available from the corresponding author upon reasonable request.

## SUPPLEMENTARTY MATERIAL

**Supplementary Figure 1.**
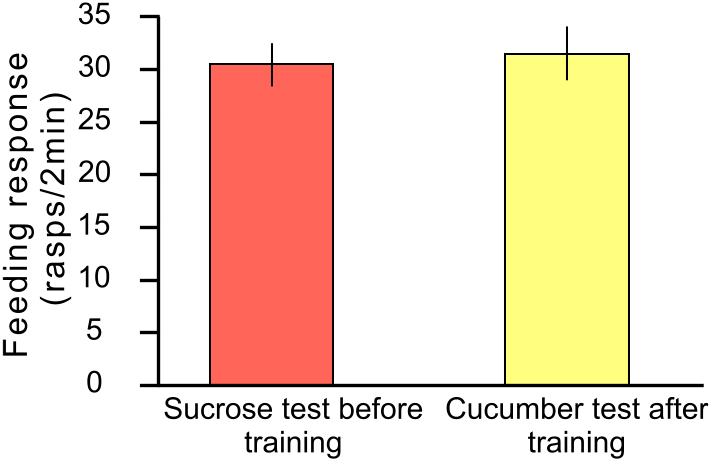
The conditioned suppression of the feeding response by aversive conditioning with sucrose as the CS does not affect the unconditioned feeding response to a different food stimulus. Fresh cucumber juice, a salient feeding stimulus, was applied 24 h after aversive training with sucrose as the CS and strong tactile stimulation as the US. The aversively conditioned animals showed the same high levels of feeding response to the cucumber juice as they had to sucrose before training.

## Notes

#### Summary of Updates

Following advice from Dr Riccardo Mozzachiodi, Texas A&M University-Corpus Christi, we now have included in the Discussion a comparison between our findings and earlier findings from a paper by Walters et al. (1981) Science, 211:4481, 504-506 that first demonstrated conditioned fear in an invertebrate, Aplysia californica. In this new discussion section we state that our findings have some interesting similarities to early seminal findings in Aplysia that provided the first evidence for conditioned fear in an invertebrate. In that study, an initially neutral chemosensory CS was paired with electric shock to the head as the US. This aversive classical conditioning resulted both in the facilitation of a number of defensive responses to the CS and a concomitant depression of the feeding response to seaweed, a natural salient food stimulus for Aplysia. However, our aversive classical conditioning paradigm was based on the use of a salient food chemosensory CS, and its primary effect was a pairing-specific depression of the feeding response to this CS, in the absence of a defensive withdrawal response to it. At the same time, the strong tactile US used in our experiments caused a sensitized response to a mild tactile stimulus at test. Therefore, unlike the paradigm used in the Aplysia study, which resulted in a variety of different conditioned fear responses to the CS, including feeding depression, our paradigm seems to have resulted in a conditioned suppression of feeding and a non-associative enhancement of the withdrawal response without the emergence of conditioned fear. From this we conclude that unlike the noxious electrical stimulation used in Aplysia, the aversive tactile stimulation used in our experiments did not produce defensive arousal.

